# Transcriptional regulation of the rainbow trout spleen corticotropin-releasing factor system in response to inflammatory challenges: roles of NF-κB and cortisol

**DOI:** 10.64898/2026.06.12.731886

**Authors:** Brett M. Culbert, Leah Grosman, Tania Rodríguez-Ramos, Brian Dixon, Nicholas J. Bernier

## Abstract

The corticotropin-releasing factor (CRF) system bidirectionally interacts with cytokines and other immune-related components in mammals. However, the nature of these interactions remains poorly characterized in other vertebrates, including teleost fishes. To gain insight into the relationship between immune responses and the CRF system in teleosts, we explored how CRF system components were transcriptionally regulated in immune organs of rainbow trout (*Oncorhynchus mykiss*). We first characterized the CRF system in the spleen and head kidney—two primary immune organs in teleosts—and found that many CRF system components were present in both tissues, but splenic expression was consistently greater. Changes in the abundance of splenic CRF system components following vaccination (which transiently stimulated inflammatory responses and cytokine production) indicated contrasting and time-dependent regulation of CRF receptor 1 (CRFR1; suppression) and CRFR2 (stimulation) activities in response to an inflammatory challenge. Using spleen explant cultures, we then evaluated whether these effects were mediated by either of nuclear factor kappa B (NF-κB; a pro-inflammatory transcription factor) or cortisol (an anti-inflammatory hormone). At baseline, cultured spleens increased cytokine production and exhibited transcriptional changes in CRF system components comparable to those observed following vaccination. Cortisol treatment and NF-κB inhibition both attenuated the rise in cytokine transcription; however, cortisol treatment generally affected transcripts influencing CRFR1 activity, while NF-κB inhibition reduced CRFR2 activity. Overall, our data provide novel insight into CRF system regulation in the spleen and suggest that cortisol and inflammatory cytokines differentially regulate CRFR1 and CRFR2 activity within this organ.

## 1. Introduction

Immunosuppression is one of the most notable consequences of chronic stress (McEwen and Seeman, 1999; Romero et al., 2009) and the anti-inflammatory actions of corticosteroid hormones are a key mediator of this effect (Cain and Cidlowski, 2017; Coutinho and Chapman, 2011). In mammals, increases in corticosteroid production cause concurrent reductions in the production of pro-inflammatory cytokines [e.g., interleukin 1β (IL-1β), interleukin 6 (IL-6), interferon γ (IFNγ), and tumour necrosis factor α (TNFa)] combined with elevated production of anti-inflammatory cytokines [e.g., interleukin 10 (IL-10)]. These effects are primarily mediated by activation of the glucocorticoid receptor, which exerts anti-inflammatory actions both via direct genomic effects (i.e., binding to glucocorticoid response elements) and transrepression of pro-inflammatory transcription factors, including nuclear factor κB (NF-κB) and activator protein-1 (Cain and Cidlowski, 2017; Coutinho and Chapman, 2011). However, these effects are not unidirectional as many immune-related factors, especially cytokines [e.g., IL-1β, IL-6, and TNFa; (Turnbull and Rivier, 1999)], can centrally regulate the hypothalamic-pituitary-adrenal/interrenal (HPA/HPI) axis and alter rates of corticosteroid synthesis. Similar bidirectional communication between stress-related hormones and immune factors has also been observed in non-mammalian vertebrates, including teleost fishes. For example, cortisol production in rainbow trout (*Oncorhynchus mykiss*) increases following injection with IL-1β (Holland et al., 2002), which then feeds back to suppress IL-1β synthesis (Zou et al., 2000). Yet, the mechanisms underlying interactions between other (non-corticosteroid) stress-related hormones and immune factors are comparatively less well understood, especially in non-mammalian vertebrates. This is especially true when considering immunoregulatory contributions unrelated to HPA/HPI axis regulation.

The corticotropin-releasing factor (CRF) system is best known for its conserved role in centrally regulating corticosteroid production (Aguilera, 1998; Best et al., 2024; Culbert & Bernier, 2026), but it also serves important functions outside of the brain. For instance, various immune cells express one or both CRF receptors (CRFR1/CRFR2), which can mediate cell- and tissue-specific pro- and anti-inflammatory effects in peripheral tissues (Dermitzaki et al. 2018; Zhu et al. 2011). Activation of CRFRs can trigger or suppress the release of inflammatory factors—such as IL-1β, IL6, and TNFa (Agelaki et al. 2002; Fleisher-Berkovich et al. 1998; Kohno et al. 2001; Leu and Singh, 1992; Torricelli et al. 2009)—and can promote survival or induce apoptosis of immune cells (Chandras et al. 2009; Harlé et al. 2018; Tsatsanis et al. 2005). There is also evidence that the CRF system has similar immunoregulatory functions in other taxa. For example, Singh and Rai (2011) reported that CRF system activation reduced phagocytosis by splenic phagocytes in spotted snakehead (*Channa punctatus*). Conversely, CRF system activation stimulated phagocytosis and promoted an anti-inflammatory phenotype (i.e., reduced production of pro-inflammatory cytokines alongside increased production of anti-inflammatory cytokines) in head kidney monocyte/macrophages of ayu [*Plecoglossus altivelis* (Jiang et al., 2021)]. In larval zebrafish (*Danio rerio*), exogenous CRF enhanced macrophage migration towards a wound site through CRFR1 and caused a modest increase in the production of some pro-inflammatory cytokines [TNFa and IL-1β, but not IL6; (van Heijningen et al., 2025)]. These observations suggest that CRF-related peptides exert diverse modulatory effects on the inflammatory response across vertebrates. Less well understood are the mechanisms by which CRF system components in immune tissues/cells are regulated. While interleukins and LPS can upregulate CRFR2 (Papadopoulou et al. 2005) in human mast cells, and LPS increases the expression of CRF and CRF receptors in mouse splenocytes (Smith et al. 2006), inflammatory challenges affect the expression CRF system components in the immune organs of non-mammalian vertebrates, such as teleost fishes, is less clear. Jiang et al. (2021) reported increased transcription of urotensin/urocortin 1 (UTS/UCN1) in the gills, head kidney, liver, and spleen following vaccination of ayu, but whether other components of the CRF system are similarly affected is unknown.

In the current study, we explored how the CRF system is transcriptionally regulated in immune tissues of rainbow trout (*Oncorhynchus mykiss*). We used rainbow trout because the regulation of immune responses (Frenette et al., 2023; Khansari et al., 2018; Soto-Dávila et al., 2024) and the peripheral CRF system [i.e., physiological roles outside of the brain (Culbert et al., 2025a; Culbert et al., 2025c; Mimassi et al., 2000)] have both been well studied in salmonids. Like most teleosts, the rainbow trout CRF system consists of five ligands [CRFa, CRFb, urotensin 1 (UTS1), urocortin 2 (UCN2), and UCN3], two receptors (CRFR1 and CRFR2), and a binding protein [CRFBP; (Cardoso et al., 2014; Cardoso et al., 2016; Maugars et al., 2022; Culbert & Bernier, 2026)], but possesses additional duplicates of all components owing to the salmonid-specific genome duplication (Berthelot et al., 2014). After first characterizing the spleen CRF system (as well as the head kidney CRF system), we then evaluated how splenic levels of the most abundant CRF system components changed following vaccination (which evoked a strong immune response). Following this, we used spleen explant cultures to determine whether the observed changes in CRF system activity were primarily mediated by systemic (factors produced outside of the spleen) or local (factors produced within the spleen) responses. Finally, we evaluated whether the observed changes in CRF system activity of spleen explants were influenced by either NF-κB or corticosteroid signalling pathways. Collectively, these experiments greatly enhance our understanding of immune-endocrine interactions in non-mammalian vertebrates.

## 2. Materials and Methods

### 2.1. Experimental Animals and Housing

All trout were acquired from the Ontario Aquaculture Research Centre (Alma, ON, Canada) and were housed in the Hagen Aqualab at the University of Guelph (Guelph, ON, Canada). Fish were initially maintained in 1.8m diameter fibreglass tanks (∼2000L) supplied with flow-through well water maintained at 12°C and kept on a 12h light, 12h dark photoperiod regime. Fish were fed to satiation three times per week with commercial trout food (3 or 5 PT Sinking; Blue Water Fish Food, Guelph, ON, Canada). A stocking density of ∼100 fish per tank was maintained during this period, and fish were kept under these conditions for at least a month prior to experiments. All procedures were carried out in accordance with the Canadian Council for Animal Care guidelines for the use of animals in research and teaching and were approved by the University of Guelph’s Animal Care Committee (AUP #4996).

### 2.2. Experiment 1: Characterization of CRF System in the Head Kidney and Spleen

To determine which components of the CRF system were expressed in the head kidney and spleen, we conducted qPCR (see Section 2.5) using tissues collected from three trout (mass= 156 ± 10 g, fork length (FL) = 23.3 ± 0.9 cm; Mean ± SEM). Values for each transcript were corrected for primer efficiency and expressed relative to the abundance of *ucn3* in the head kidney, the CRF system component with the lowest levels across both tissues that consistently amplified. We measured individual paralogs of all CRF system components, except for UCN3, which shares 99% identity across paralogs.

### 2.3. Experiment 2: Regulation of the Spleen CRF System following Vaccination

Trout (N = 94; mass = 270 ± 6 g, FL = 27.7 ± 0.2 cm) were lightly anaesthetized using MS-222 (70 mg L^-1^; Syndel, Nanaimo, BC, Canada), weighed, and transferred into one of nine experimental tanks (N=9-11 fish per tank). Fish were held in 0.6 m diameter fibreglass tanks (∼200L) supplied with flow-through well water and containing an air stone. All tanks were maintained at 12°C and kept on a 12h light, 12h dark photoperiod regime. Trout were fed 1% of the average body weight in each tank daily and were held under these conditions for two weeks. Following this acclimation period, fish (N=10) in one tank were sampled as a control group. The remaining fish were anesthetized using MS-222 (100 mg L^-1^) and injected intraperitoneally with 250 µL of either an inactivated vaccine solution or filter-sterilized saline (0.9% NaCl). Food was withheld for 24 h prior to injection. The vaccine (ICTHIOVAC VR; HIPRA, Amer, Girona, Spain) contained formalin-killed *Vibrio anguillarum* (serotypes O1, O2α and O2β) and has previously been shown to elicit an inflammatory response in rainbow trout (Khansari et al., 2018; Khansari et al., 2019; Liu et al., 2019). After either 6, 24, 72, or 168 h post-injection, groups of vaccine- and saline-injected fish were terminally anesthetized using 2-phenoxyethanol (0.2%; Sigma-Aldrich, Oakville, ON, Canada). Fork length and mass were then recorded, after which blood was collected from the caudal vasculature using a 1 mL ammonium heparinized syringe. Blood was centrifuged at 9,000 *g* for 5 min, and plasma was collected and frozen on dry ice. Afterwards, the spleen was removed and frozen on dry ice. Plasma and spleen were subsequently stored at -80°C to be used for later evaluation of cortisol levels (plasma; see Section 2.5) or protein/transcript levels (spleen; see Sections 2.6 and 2.7). Specifically, we measured protein levels of two cytokines (IL-1β and IFNγ), as well as transcript abundance of cytokines (*il1b*, *il6*, *tnfa*, *ifny*, and *il10*) and the major components of the splenic CRF system (*crfa1*, *ucn3*, *uts1a*,*crfbp1, crfr1a, crfr1b, crfr2a*, and *crfr2b*; see Fig. 1). All fish were sampled at 1400 h and food was withheld throughout the experiment to avoid potential confounds associated with the inhibition of appetite due to vaccination (Bernier, 2010; Sørum and Damsgård, 2004).

**Figure 1.**
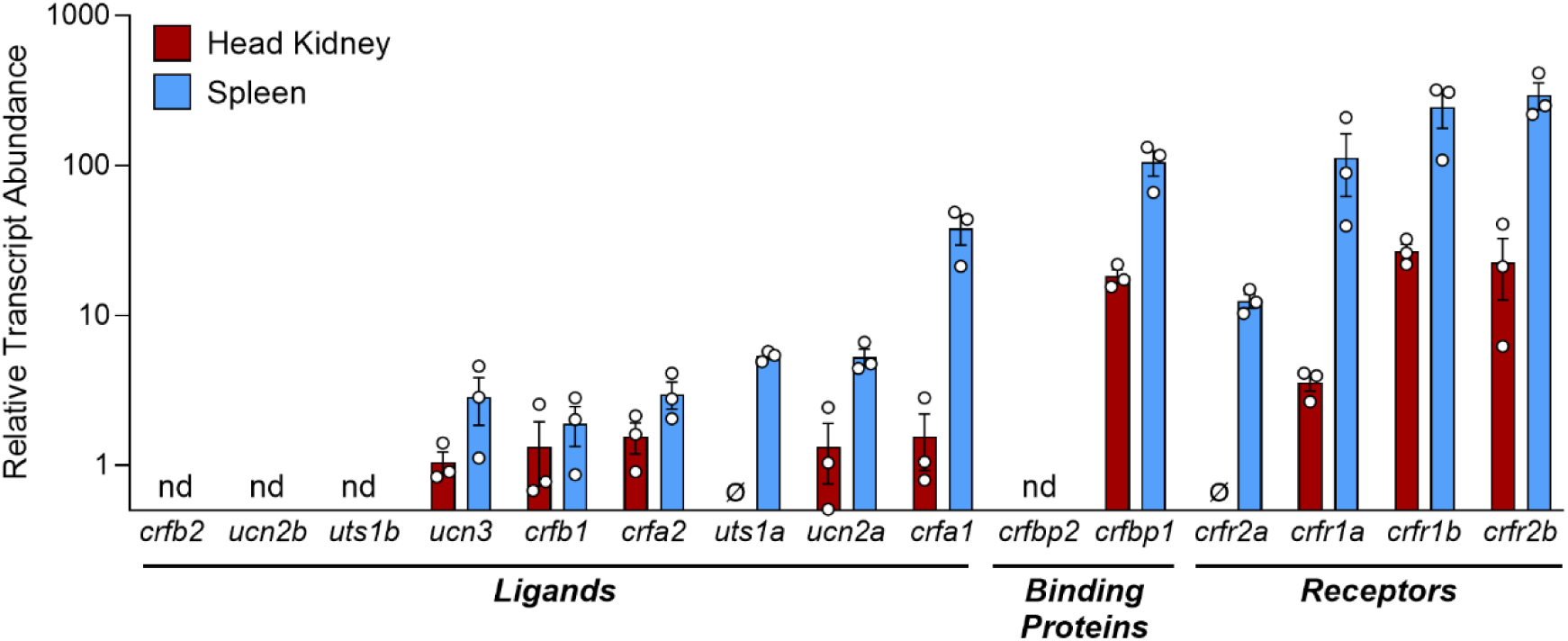
Relative abundance of individual ligands, binding proteins, and receptors of the corticotropin-releasing factor (CRF) system in the head kidney (red) and spleen (blue) of rainbow trout (*Oncorhynchus mykiss*). Bars represent mean abundance (±SEM) of each component relative to *ucn3* in the head kidney (the component with the lowest detectable levels). Each point represents an individual value from an unstressed fish. Components that were not detected in any of the three tissues are indicated with nd (not detected) and the Ø symbol indicates instances where components were only undetectable in the spleen. Note that data are plotted on a Log10 scale for visualization purposes. *crf*, corticotropin-releasing factor; *crfbp*, corticotropin-releasing factor binding protein; *crfr*, corticotropin-releasing factor receptor; *ucn*, urocortin; *uts*, urotensin.

### 2.4. Experiment 3: *In vitro* Regulation of the CRF System in Spleen Explants

#### 2.4.1. Effects of an Endogenous Inflammatory Response on the CRF System

Trout (N = 8; mass = 662 ± 59 g, FL = 36.6 ± 1.2 cm) were terminally anaesthetized using 2-phenoxyethanol (0.2%) and fork length and mass were recorded. Afterwards, the spleen was removed and split into three pieces. One piece was immediately frozen on dry ice as a control, while the other two pieces were placed into wells of a 24 well plate containing 0.75 mL of minimum essential medium with Hanks’ Balanced salts (MEM; GIBCO, Product #11575032) that was supplemented with 500 U mL^-1^ penicillin/streptomycin (GIBCO, Product #15140122), 4 mg mL^-1^ bovine serum albumin (BSA), and 25 mM HEPES (final pH of 7.55). These pieces were incubated with shaking at 12°C for either 6 or 24 h, after which they were collected and frozen on dry ice to facilitate later RNA extraction and qPCR analysis (see Section 2.6). We measured the same transcripts that were measured in Experiment 2.

#### 2.4.2. Effects of NF-κB Signalling Blockade on the CRF System

Trout (N = 8; mass = 211 ± 16 g, FL = 26.4 ± 0.6 cm) were processed as above (Section 2.4.1), with the exception that spleen pieces were incubated in MEM that contained either vehicle (VEH; MEM with 0.1% ddH_2_O) or the NF-κB inhibitor pyrrolidine dithiocarbamate (PDTC; MEM with 0.1% ddH_2_O and 100 µM of PDTC) for 6 and 24 h. Previous work has shown that this dose is sufficient to attenuate *in vitro* cytokine production across a variety of tissues and species (Cao et al., 2017; Munoz et al., 1996; Rangan et al., 1999; Zhou et al., 2022). After this incubation period, spleen pieces were collected, flash frozen on dry ice, and stored at - 80°C to facilitate later RNA extraction and qPCR analysis (see Section 2.6). We measured the same transcripts that were measured in Experiment 2. Pyrrolidine-1-dithiocarboxylic acid ammonium salt (Product # 821100) was purchased from Sigma-Aldrich.

#### 2.4.3. Effects of Cortisol and Glucocorticoid Receptor Signalling on the CRF System

Trout (N = 8; mass = 778 ± 108 g, FL = 37.0 ± 2.3 cm) were processed as above (Section 2.4.1), with the exception that spleen pieces were incubated in MEM that contained either vehicle alone (VEH; MEM with 0.1% EtOH), cortisol alone (CORT; MEM with 0.1% EtOH and 10 ug mL^-1^ of cortisol), or a combination of cortisol and the glucocorticoid receptor antagonist RU486 (CORT+RU486; MEM with 0.1% EtOH, 10 ug mL^-1^ of cortisol, and 10 ug mL^-1^ of RU486) for 6 and 24 h. Previous work has shown that these doses are sufficient to manipulate corticosteroid receptor activity *in vitro* in salmonids (Culbert et al., 2025a; Kiilerich et al., 2007; Pagniello et al., 2002; Tipsmark et al., 2009). Afterwards, spleen pieces were collected, flash frozen on dry ice, and stored at -80°C to facilitate later RNA extraction and qPCR analysis (see Section 2.6). We measured the transcripts that were measured in Experiment 2. Cortisol (Product # H4001) and RU486 (Product # M8046) were both purchased from Sigma-Aldrich.

### 2.5. Determination of Plasma Cortisol Concentrations

Plasma cortisol levels were measured in duplicate using a previously validated, commercially available enzyme-linked immunosorbent assay (ELISA; Cat # 402710; Neogen, Lexington, KY, USA). Intra- and inter-assay coefficients of variation were 9.3% and 4.5%, respectively.

### 2.6. Determination of Cytokine Protein Levels in the Spleen

Protein levels of IL-1β (5.7 and 13.0 % CV, intra- and inter-assay) and IFNγ (3.4 and 14.6 % CV, intra- and inter-assay) were measured in triplicate using previously validated salmonid-specific sandwich ELISAs (see Frenette et al., 2023 and Soto-Dávila et al., 2024 for more details). To extract protein from the spleen, ∼100 mg of tissue was homogenized in T-PER containing protease inhibitors (Thermo Fisher Scientific, Mississauga, ON, Canada) and centrifuged at 10,000 *g* for 5 mins. The supernatant was collected and total protein concentration was determined using a commercial bicinchoninic acid assay (Thermo Fisher Scientific). For both ELISAs, 50 µg of protein was added to each well.

### 2.7. RNA Isolation and qPCR

Spleens were homogenized in TRIzol reagent (Invitrogen, Burlington, ON, Canada) using a Precellys Evolution tissue homogenizer (Bertin Instruments, Montigny-le-Bretonneux, France). Following the manufacturer’s protocol, total RNA was extracted, and its quantity and purity were assessed using a NanoDrop 2000 spectrophotometer (Thermo Fisher Scientific). Following this, we treated 1 µg of RNA with DNase (DNase 1; Thermo Fisher Scientific) and reverse transcribed cDNA using a high-capacity cDNA reverse transcription kit (Applied Biosystems, Waltham, MA, USA). We then performed qPCR using a CFX96 system (BioRad, Hercules, CA, USA) with SYBR green (SsoAdvanced Universal; BioRad) and gene-specific primers (Supp. Table 1). All samples were run in duplicate and negative controls—including no template controls (where cDNA was replaced with water) and no reverse transcriptase controls (where reverse transcriptase was replaced with water during cDNA synthesis)—were also included. Each reaction contained a total of 20 µl, which consisted of 10 µl of SYBR green, 5 µl of combined forward and reverse primers (0.2 µM [final]), and 5 µl of 10x diluted cDNA. Cycling parameters included a 30 s activation step at 95°C, followed by 40 cycles consisting of a 3 s denaturation step at 95°C and a combined 30 s annealing and extension step at 60°C. Melt curve analysis was conducted at the end of each run to confirm the specificity of each reaction. To account for differences in amplification efficiency, standard curves were constructed for each gene using serial dilutions (4x) of pooled cDNA. Input values for each gene were obtained by fitting the average threshold cycle value to the antilog of the gene-specific standard curve, thereby correcting for differences in primer amplification efficiency. To correct for minor variations in template input and transcriptional efficiency, we normalized our data to the geometric mean of transcript abundances of elongation factor 1α (*ef1α*) and ribosomal protein L13a (*rpl13a*), which served as reference genes (both were stable across groups in all experiments). All data are expressed relative to the mean value of the control group within each experiment (see figure captions for further details).

**Table 1.** Summary of transcriptional responses of the major splenic corticotropin-releasing factor (CRF) system components observed across all experiments. Note that effects of NF-κB signalling are inferred based on responses following NF-κB inhibition. crfbp, CRF binding protein; crfr, CRF receptor; NC, no change; NF-κB, nuclear factor κB; ucn, urocortin; uts, urotensin.

|  | Vaccination | Explant Protocol | NF-κB Signalling | Cortisol Signalling |
| --- | --- | --- | --- | --- |
| *crfr2b* | ↑ | ↑ | ↑ | ↑ |
| *crfr2a* | ↑ | ↓ | NC | NC |
| *crfr1b* | ↓ | ↓ | NC | ↑ |
| *crfr1a* | NC | NC | NC | ↓ |
| *crfbp1* | ↑ | ↑ | NC | ↓ |
| *crfa1* | ↓ | ↓ | NC | ↓ |
| *uts1a* | ↓ | ↑ | NC | ↑ |
| *ucn3* | ↑ | ↑ | ↑ | NC |

### 2.8. Statistical Analyses

Statistical analyses were performed using R [v4.4.0; (R Core Team, 2026)]. All data are presented as means ± 1 standard error of the mean (SEM), and a significance level (α) of 0.05 was used for all tests. Outliers were excluded based on a 2x interquartile range threshold. When data did not meet the assumptions of normality and/or equal variance, they were log or square-root transformed to improve fit. Data for Experiment 2 were analyzed using two-way ANOVAs that included treatment (saline- and vaccine-injected) and time post-injection (6, 24, 72, and 168 h), as well as the interaction between these factors. Data for Experiment 3 were either analyzed using one-way ANOVAs (culture effects; 0, 6, and 24 h) or two-way ANOVAs that included treatment (cortisol effects: VEH, CORT, and CORT+RU486; NF-κB effects: VEH and PDTC) and culture duration (6 and 24 h), as well as the interaction between these factors. Since pieces of spleen from the same fish were included in all treatment groups for trials in Experiment 3, we included fish ID as a random effect using the ‘lme4’ package (Bates et al., 2015). When significant differences were detected, post hoc Tukey’s tests were performed using the ‘emmeans’ package (Lenth, 2016).

## 3. Results

### Abundance of CRF System Components in the Head Kidney and Spleen

The majority of CRF system components were reliably detected in both the head kidney and spleen (Fig. 1); however, their abundance was consistently higher in the spleen than the head kidney. All CRF ligands were detectable except for *crfb2* (undetectable in both tissues), *ucn2b* (undetectable in both tissues), *uts1b* (undetectable in both tissues), and *uts1a* (undetectable in head kidney). Of the CRF ligands which were detected, splenic *crfa1* displayed the highest abundance (amplified at ∼30.5 cycles), and was ∼7-20x and ∼25-35x greater than other ligands in the spleen and head kidney, respectively. For binding proteins, only *crfbp1* was detectable and was ∼5x higher in the spleen compared to the head kidney. Finally, all CRF receptors were detectable except for *crfr2a* in the head kidney. Abundance of *crfr2b* (spleen: ∼27.5 cycles; head kidney: ∼31.5 cycles) and *crfr1b* (spleen: ∼28.5 cycles; head kidney: ∼31 cycles) was the greatest across both tissues, but levels in the spleen were ∼9x and ∼13x higher than in the head kidney, respectively.

### Effects of Vaccination on Splenic Cytokines, Plasma Cortisol, and CRF System Components

Splenic levels of IL-1β were elevated at the transcript (Fig. 2A; p_treatment_<0.001, p_time_<0.001, p_treatment*time_<0.001) and protein levels (Fig. 2B; p_treatment_<0.001, p_time_<0.001, p_treatment*time_<0.001) following vaccination, with peak responses of ∼3700- and 5-fold, respectively, occurring 6 h post-vaccination. Transcript levels of *il6* (Fig. 2C; p_treatment_<0.001, p_time_<0.001, p_treatment*time_<0.001) and *tnfa* (Fig. 2D; p_treatment_<0.001, p_time_<0.001, p_treatment*time_<0.001) also peaked at 6 h post-vaccination, displaying increases of 1200- and 65-fold, respectively. In contrast, maximal transcriptional responses of *ifny* (Fig. 2E; p_treatment_<0.001,p_time_<0.001, p_treatment*time_<0.001) were not observed until 24 h post-vaccination (increasing ∼30-fold), and protein levels of IFNγ (Fig. 2F; p_treatment_=0.003, p_time_=0.52, p_treatment*time_=0.84) were ∼2-fold higher in vaccinated fish across all timepoints. Peak transcript abundance of *il10* (Fig. 2F; p_treatment_<0.001, p_time_<0.001, p_treatment*time_<0.001) was also not observed until 24 h post-vaccination, displaying an increase of ∼700-fold. For all cytokines examined, no differences were observed following saline injection and levels in vaccinated fish had generally returned to levels consistent with saline-injected fish by 168 h post-vaccination. Additionally, circulating cortisol levels (Fig. 2H; p_treatment_=0.01, p_time_<0.001, p_treatment*time_=0.06) were ∼30% higher overall in vaccinated fish compared to saline-injected fish and were also elevated 2‒4-fold overall at 6 h post-injection compared to all other timepoints.

**Figure 2.**
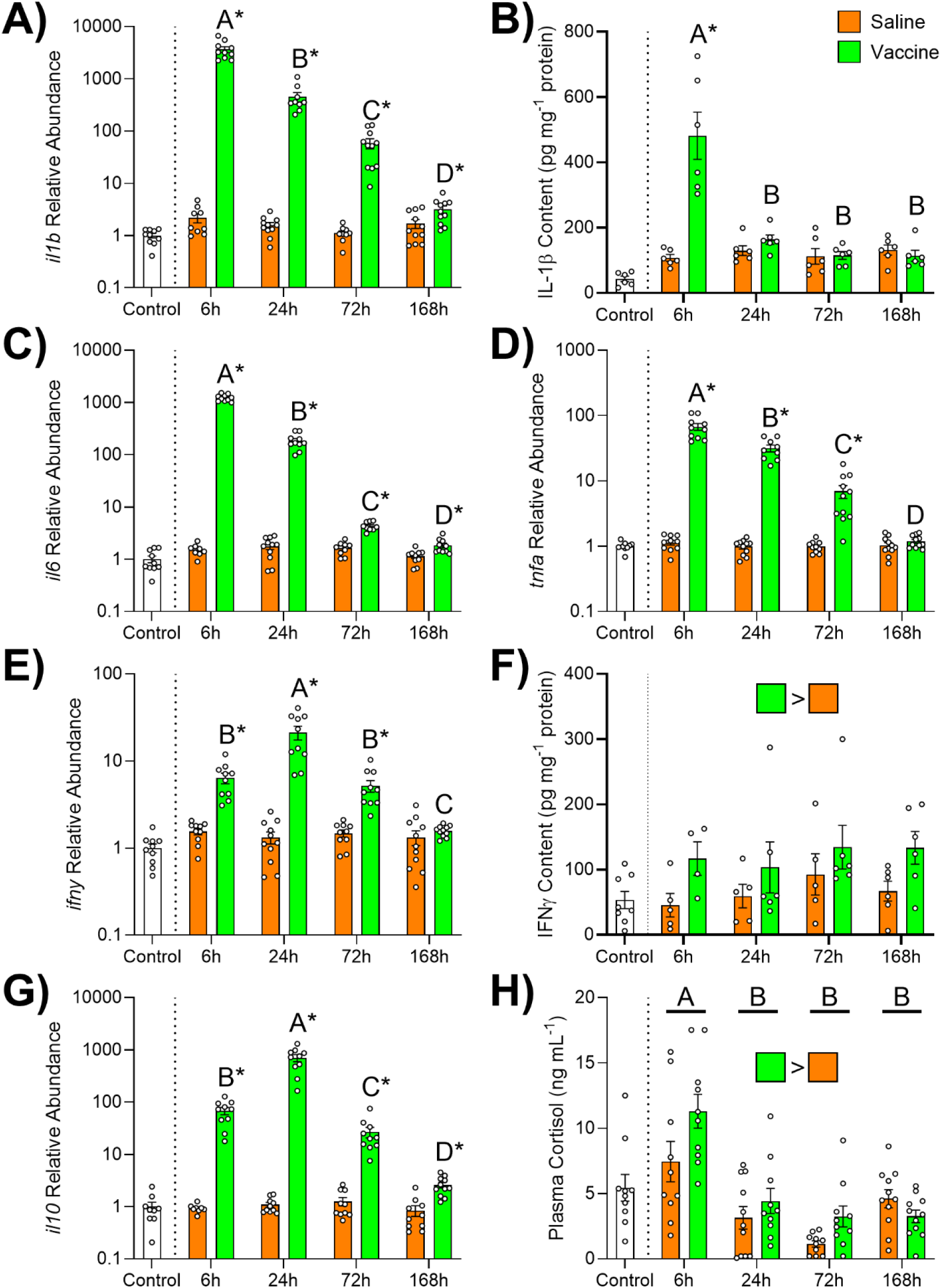
Time-dependent effects of saline (orange) or vaccine (green) injection on splenic interleukin 1β transcript (*il1b*, A) and protein abundance (IL-1β, B), transcript abundance of interleukin 6 (*il6*; C) and tumor necrosis factor α (*tnfa*; D), interferon γ transcript (*ifny*, E) and protein abundance (IFNγ, F), transcript abundance of interleukin 10 (*il10*; G), and plasma cortisol levels (H), in rainbow trout (*Oncorhynchus mykiss*). Significant differences (p < 0.05; two-way ANOVA) are depicted using either letters (across time; uppercase = within vaccine group), filled oversized squares (between groups across all timepoints) or asterisks (between groups within a timepoint). Transcript abundance data are expressed relative to the control group (white), but control fish were not included in any analyses. Note that transcript abundance data are plotted on a Log10 scale for visualization purposes. Values are represented as means ± SEM and individual data points are shown.

Transcript levels of splenic *crfr2b* (Fig. 3A; p_treatment_<0.001, p_time_<0.001, p_treatment*time_<0.001) and *crfr2a* (Fig. 3B; p_treatment_=0.05, p_time_=0.002, p_treatment*time_=0.005) showed similar responses following vaccination, whereby levels were ∼3-fold higher 24 h post-vaccination. In contrast, levels of *crfr1b* (Fig. 3C; p_treatment_<0.001, p_time_=0.04, p_treatment*time_<0.001) were ∼85% lower 24 h post-vaccination, with a similar (but non-significant) pattern for *crfr1a* (Fig. 3D; p_treatment_=0.16, p_time_=0.08, p_treatment*time_=0.23). Abundance of *crfbp1* (Fig. 3E; p_treatment_<0.001, p_time_<0.001, p_treatment*time_<0.001) was 2-2.5x higher at 6 and 24 h post-vaccination, but values in vaccinated fish did not differ from saline-injected fish at 72 or 168 h. Levels of *crfa1* (Fig. 3F; p_treatment_<0.001, p_time_=0.002, p_treatment*time_=0.73) and *uts1a* (Fig. 3G; p_treatment_<0.001, p_time_<0.001, p_treatment*time_=0.19) were both lower overall in vaccinated fish compared to saline-injected controls. Additionally, *crfa1* was lower in both groups at 24 h post-injection, and *uts1a* was lower in both groups at 6 h post-injection. In contrast, levels of *ucn3* (Fig. 3H; p_treatment_=0.49, p_time_<0.001, p_treatment*time_<0.001) increased within 72 h in both groups, but this increase occurred earlier (by 6 h post-injection) and returned to baseline more quickly (by 72 h post-injection) in vaccinated fish.

**Figure 3.**
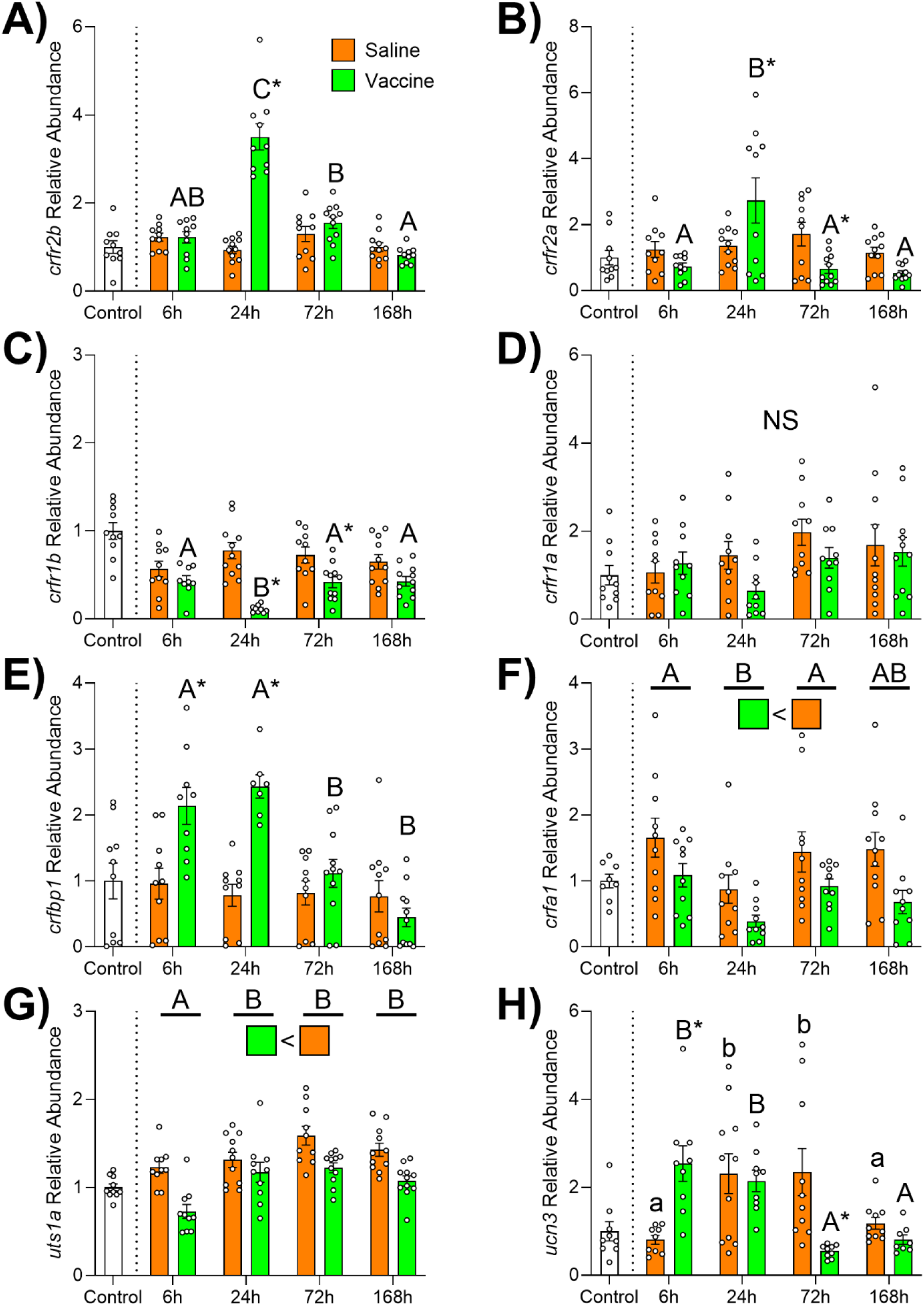
Time-dependent effects of saline (orange) or vaccine (green) injection on transcript abundance of corticotropin-releasing factor (CRF) receptor 2b (*crfr2b*; A), CRF receptor 2a (*crfr2a*; B), CRF receptor 1b (*crfr1b*; C), CRF receptor 1a (*crfr1a*; D), CRF binding protein 1 (*crfbp1*; E), CRFa1 (*crfa1*; F), urotensin 1a (*uts1a*; G), and urocortin 3 (*ucn3*; H) in the spleen of rainbow trout (*Oncorhynchus mykiss*). Significant differences (p < 0.05; two-way ANOVA) are depicted using either letters (across time; uppercase = within vaccine group, underlined uppercase = overall time effect), filled oversized squares (between groups across all timepoints) or asterisks (between groups within a timepoint). Data are expressed relative to the control group (white), but control fish were not included in the analyses. Values are represented as means ± SEM and individual data points are shown. NS: nonsignificant.

### Changes in Cytokine and CRF System Component Levels in Spleen Explants

Levels of *il1b* (Fig. 4A; p_time_<0.001) and *il6* (Fig. 4A; p_time_<0.001) transcripts were both several hundred-fold higher after 6 or 24 h of culture compared to baseline. A time-dependent response was observed for *tnfa* (Fig. 4A; p_time_<0.001), with levels being ∼5 and ∼25x higher than baseline at 6 h and 24 h, respectively; however, while values at 24 h were significantly different from 0 h (p<0.001), values at 6 h were not (p=0.06). Effects on *ifny* (Fig. 4A; p_time_=0.002) and *il10* (Fig. 4A; p_time_<0.001) were consistent at both 6 h and 24 h, although, the direction of these effects differed. While *ifny* levels were reduced by ∼40%, *il10* abundance was ∼15-fold higher.

**Figure 4.**
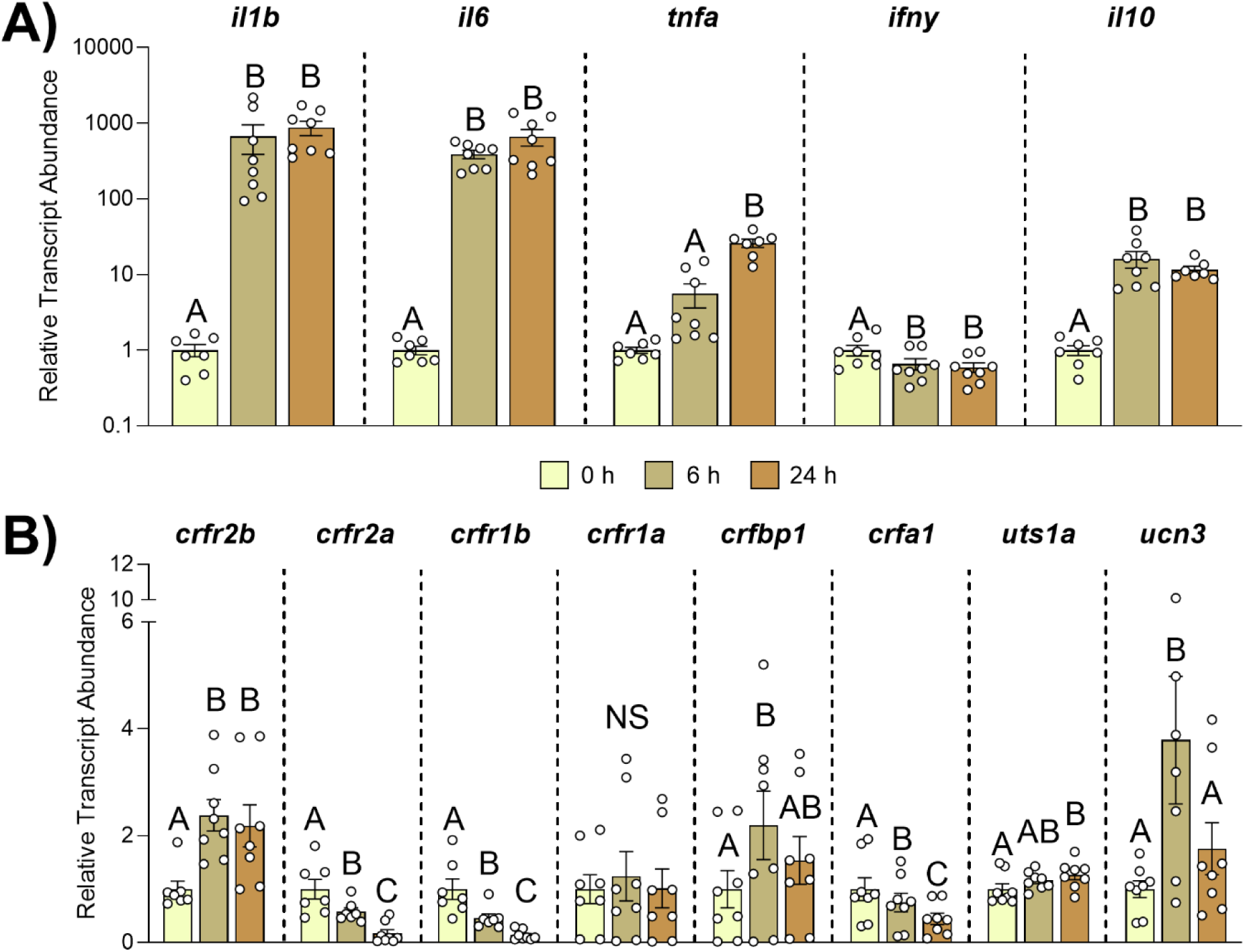
Time-dependent effects *in vitro* culture on transcript abundance of immune-related cytokines [A; interleukin 1β (*il1b*), interleukin 6 (*il6*), tumor necrosis factor α (*tnfa*), interferon γ (*ifny*), and interleukin 10 (*il10*)] and components of the corticotropin-releasing factor (CRF) system [B; CRF receptor 2b (*crfr2b*), CRF receptor 2a (*crfr2a*), CRF receptor 1b (*crfr1b*), CRF receptor 1a (*crfr1a*), CRF binding protein 1 (*crfbp1*), CRFa1 (*crfa1*), urotensin 1a (*uts1a*), and urocortin 3 (*ucn3*)] in the spleen of rainbow trout (*Oncorhynchus mykiss*). Spleens that were cultured for 0, 6, and 24 h are depicted in beige, brown and chestnut, respectively. Significant differences (p < 0.05; one-way ANOVA) are depicted using letters. Data are expressed relative to the 0 h group. Note that data in Panel A are plotted on a Log10 scale for visualization purposes. Values are represented as means ± SEM and individual data points are shown. NS: nonsignificant.

The abundance of *crfr2b* (Fig. 4B; p_time_<0.001) transcripts was elevated ∼2.2-fold at 6 h and 24 h compared to baseline. In contrast, both *crfr2a* (Fig. 4B; p_time_<0.001) and *crfr1b* (Fig.4B; p_time_<0.001) displayed time-dependent reductions, reaching levels that were ∼15% of baseline levels by 24 h. No changes were detected for *crfr1a* (Fig. 4B; p_time_=0.22). Levels of *crfbp1* (Fig. 4B; p_time_=0.03) were ∼2.2-fold higher than baseline levels at 6 h (p=0.04) but declined to levels which were not different from either 0 h (p=0.48) or 6 h (p=0.35) after 24 h. Levels of *crfa1* (Fig. 4B; p_time_<0.001) and *uts1a* (Fig. 4B; p_time_=0.004) both displayed time-dependent changes; however, whereas levels of *crfa1* gradually declined to levels that were∼45% of baseline by 24 h, abundance of *uts1a* had increased by 27% after 24 h. Levels of *ucn3* (Fig. 4B; p_time_<0.001) increased ∼4-fold compared to baseline after 6 h (p<0.001), but returned to levels that were not different from baseline after 24 h (p=0.29).

### Effects of NF-κB Inhibition on Splenic Cytokine and CRF System Component Levels

Levels of *il1b* (Fig. 5A; p_treatment_<0.001, p_time_<0.001, p_treatment*time_=0.23) transcripts were∼50% lower overall following PDTC treatment and increased by ∼4-fold in both groups at 24 h compared to 6 h. While abundance of *il6* (Fig. 5A; p_treatment_<0.001, p_time_<0.001, p_treatment*time_<0.001) increased ∼3.5-fold after 24 h in the vehicle-treated group (p<0.001), levels did not change across time in the PDTC-treated group (p=0.63) and were ∼75% lower than the vehicle-treated group at 24 h (p<0.001). Levels of *tnfa* (Fig. 5A; p_treatment_=0.85, p_time_<0.001, p_treatment*time_=0.98) were ∼12-fold higher at 24 h compared to 6 h in both groups, but no treatment effects were observed. Levels of *ifny* (Fig. 5A; p_treatment_=0.04, p_time_=0.11, p_treatment*time_=0.56) and *il10* (Fig. 5A; p_treatment_=0.005, p_time_<0.001, p_treatment*time_=0.18) were ∼25% and ∼30% lower, respectively, in the PDTC-treated group compared to the vehicle-treated group across both timepoints. In addition, levels of *il10* were ∼55% higher at 24 h compared to 6 h across both groups.

**Figure 5.**
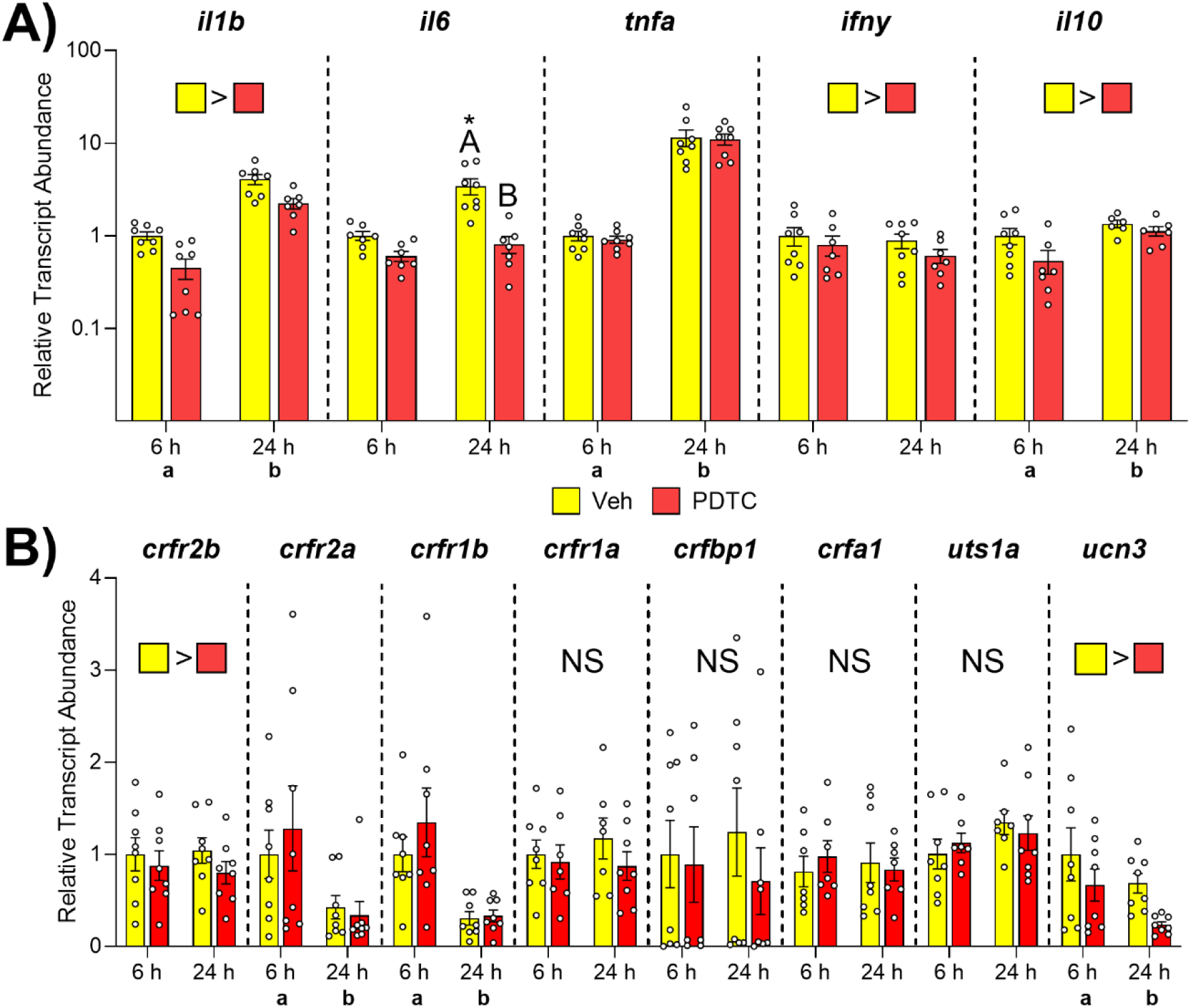
*In vitro* effects of nuclear factor κB (NF-κB) signalling blockade on transcript abundance of immune-related cytokines [A; interleukin 1β (*il1b*), interleukin 6 (*il6*), tumor necrosis factor α (*tnfa*), interferon γ (*ifny*), and interleukin 10 (*il10*)] and components of the corticotropin-releasing factor (CRF) system [B; CRF receptor 2b (*crfr2b*), CRF receptor 2a (*crfr2a*), CRF receptor 1b (*crfr1b*), CRF receptor 1a (*crfr1a*), CRF binding protein 1 (*crfbp1*), CRFa1 (*crfa1*), urotensin 1a (*uts1a*), and urocortin 3 (*ucn3*)] in the spleen of rainbow trout (*Oncorhynchus mykiss*). Spleens that were treated with vehicle (Veh) or an NF-κB antagonist (PDTC) (PDTC) are depicted in yellow and red, respectively. Significant differences (p < 0.05; two-way ANOVA) are depicted using either letters (lowercase = treatment effect at 6 h, uppercase = treatment effect at 24 h, letters along x-axis = overall time effect), filled oversized squares (treatment effect across both timepoints) or asterisks (within a group across time). Data are expressed relative to the vehicle group at 6 h. Note that data in Panel A are plotted on a Log10 scale for visualization purposes. Values are represented as means ± SEM and individual data points are shown. NS: nonsignificant.

Levels of *crfr2b* (Fig. 5B; p_treatment_=0.008, p_time_=0.79, p_treatment*time_=0.40) transcripts were∼20% lower in the PDTC-treated group across both timepoints. No treatment effects were detected for either *crfr2a* (Fig. 5B; p_treatment_=0.95, p_time_<0.001, p_treatment*time_=0.39) or *crfr1b* (Fig. 5B; p_treatment_=0.15, p_time_<0.001, p_treatment*time_=0.73), but levels of both transcripts were ∼60-70% lower in both groups at 24 h compared to 6 h. No significant effects were detected for *crfr1a* (Fig. 5B; p_treatment_=0.20, p_time_=0.78, p_treatment*time_=0.58), *crfbp1* (Fig. 5B; p_treatment_=0.13, p_time_=0.89, p_treatment*time_=0.14), *crfa1* (Fig. 5B; p_treatment_=0.42, p_time_=0.51, p_treatment*time_=0.31), or *uts1a* (Fig.5B; p_treatment_=0.84, p_time_=0.11, p_treatment*time_=0.19). However, levels of *ucn3* (Fig. 5B; p_treatment_=0.007, p_time_=0.01, p_treatment*time_=0.69) were ∼50% lower in the PDTC-treated group compared to the vehicle-treated group and were ∼50% lower in both groups at 24 h compared to 6 h.

### Effects of Cortisol Signalling on Splenic Cytokine and CRF System Component Levels

Transcript abundance of *il1b* (Fig. 6A; p_treatment_=0.02, p_time_=0.03, p_treatment*time_=0.58) was ∼50% lower overall in the cortisol-treated group compared to the vehicle-treated group (p=0.03) and also tended to be lower than the cortisol+RU486-treated group; however, this difference did not quite reach statistical significance (p=0.06). Additionally, *il1b* levels were also ∼50% higher across both groups at 24 h compared to 6 h. Levels of *il6* (Fig. 6A; p_treatment_<0.001, p_time_=0.12, p_treatment*time_=0.08) were ∼80% and ∼65% lower across both timepoints in cortisol-treated spleens compared to vehicle- (p<0.001) and cortisol+RU486-treated spleens (p<0.001), respectively. Levels of *tnfa* (Fig. 6A; p_treatment_<0.001, p_time_<0.001, p_treatment*time_<0.001) and *ifny* (Fig. 6A; p_treatment_<0.001, p_time_=0.01, p_treatment*time_=0.36) were also reduced following cortisol treatment; however, while levels in cortisol-treated spleens were only lower than vehicle- (p<0.001) and cortisol+RU486-treated spleens (p<0.001; by 95% and 85%, respectively) after 24 h for *tnfa*, levels of *ifny* were lower in cortisol-treated spleens compared to vehicle- (p<0.001) and cortisol+RU486-treated spleens across both timepoints (p<0.001; by ∼70% and ∼45%, respectively). Additionally, for both *tnfa* (only after 24 h; p=0.002) and *ifny* (overall; p<0.001), transcript levels in the cortisol+RU486-treated group remained ∼50% lower than the vehicle-treated group. Abundance of *il10* (Fig. 6A; p_treatment_<0.001, p_time_=0.006, p_treatment*time_=0.007) was elevated in the cortisol-treated group compared to the vehicle- and cortisol+RU486-treated groups at 6 h (by ∼3.5-fold; p=0.005 and p=0.006, respectively) and 24 h (by 21- and 14-fold, respectively; both p<0.001). While levels of *il10* did not change across time following vehicle (p=0.89) or cortisol+RU486 treatment (p=0.42), levels were 5x higher after 24 h compared to 6 h of cortisol treatment (p<0.001).

**Figure 6.**
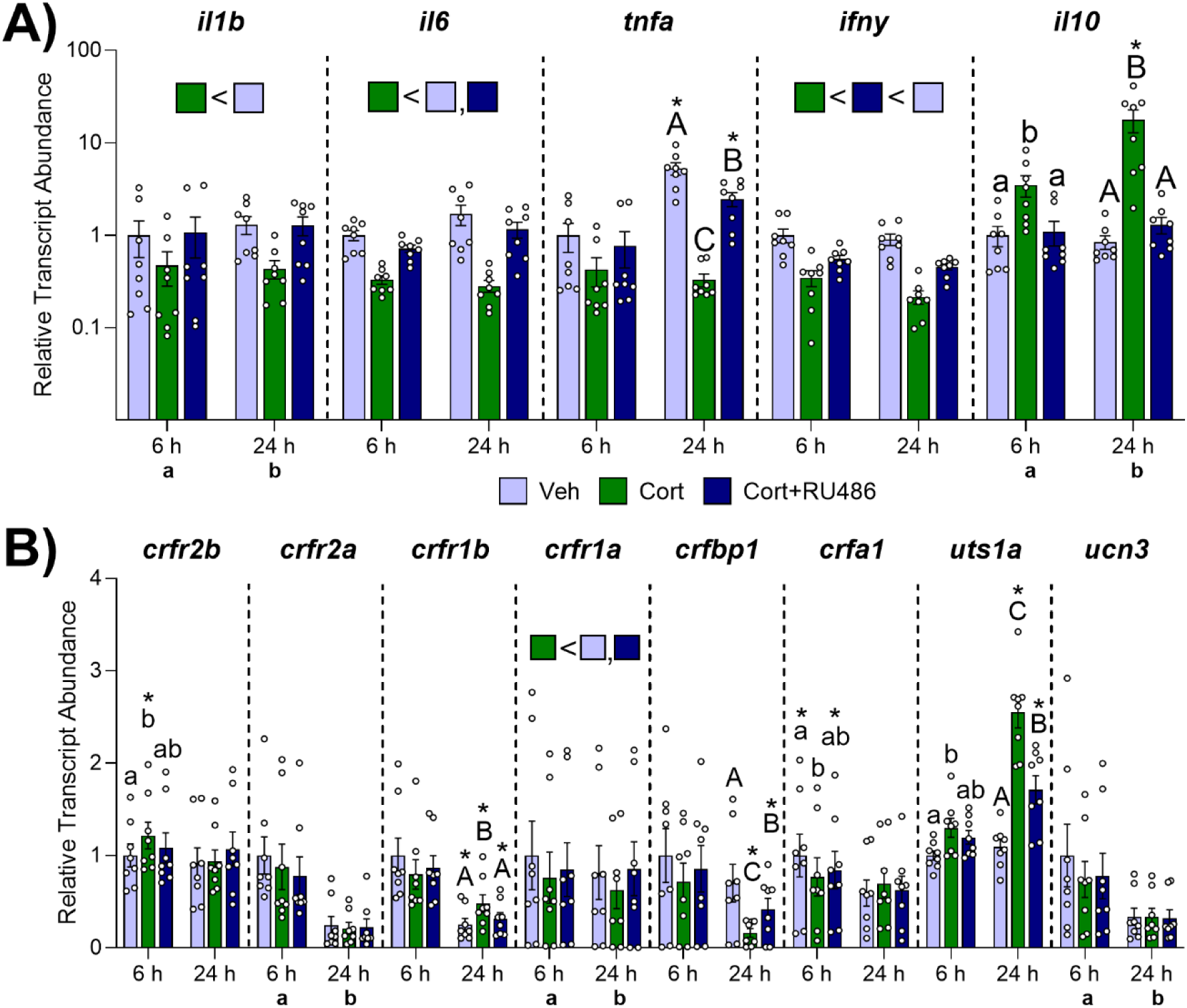
*In vitro* effects of cortisol signalling on transcript abundance of immune-related cytokines [A; interleukin 1β (*il1b*), interleukin 6 (*il6*), tumor necrosis factor α (*tnfa*), interferon γ (*ifny*), and interleukin 10 (*il10*)] and components of the corticotropin-releasing factor (CRF) system [B; CRF receptor 2b (*crfr2b*), CRF receptor 2a (*crfr2a*), CRF receptor 1b (*crfr1b*), CRF receptor 1a (*crfr1a*), CRF binding protein 1 (*crfbp1*), CRFa1 (*crfa1*), urocortin 3 (*ucn3*), and urotensin 1a (*uts1a*)] in the spleen of rainbow trout (*Oncorhynchus mykiss*). Spleens that were treated with vehicle (Veh), cortisol (Cort) or cortisol plus the glucocorticoid receptor antagonist RU486 (Cort+RU486) are depicted in light blue, green and dark blue, respectively. Significant differences (p < 0.05; two-way ANOVA) are depicted using either letters (lowercase = treatment effect at 6 h, uppercase = treatment effect at 24 h, letters along x-axis = overall time effect), filled oversized squares (treatment effect across both timepoints) or asterisks (within a group across time). Data are expressed relative to the vehicle group at 6 h. Note that data in Panel A are plotted on a Log10 scale for visualization purposes. Values are represented as means ± SEM and individual data points are shown.

Levels of *crfr2b* (Fig. 6B; p_treatment_=0.03, p_time_=0.003, p_treatment*time_=0.03) transcripts were ∼20% higher in the cortisol-treated group compared to the vehicle-treated group at 6 h (p=0.01)—but were not different from the cortisol+RU486-treated group (p=0.16)—and no differences were detected after 24 h (all p≥0.10). Abundance of *crfr2a* (Fig. 6B; p_treatment_=0.39, p_time_<0.001, p_treatment*time_=0.51) was ∼75% lower in all groups at 24 h versus 6 h, but no treatment effect was observed. Levels of *crfr1b* (Fig. 6B; p_treatment_=0.01, p_time_<0.001, p_treatment*time_<0.001) were ∼60% lower at 24 h compared to 6 h in all groups (all p<0.001), but levels were ∼90% and ∼50% higher in the cortisol-treated group (both p≤0.001) compared to the vehicle- and cortisol+RU486-treated groups at 24 h, respectively. In contrast, levels of *crfr1a* (Fig. 6B; p_treatment_=0.002, p_time_=0.049, p_treatment*time_=0.30) were ∼25% lower in the cortisol-treated group compared to the vehicle- (p=0.005) and cortisol+RU486-treated groups (p=0.048) across both timepoints. Additionally, *crfr1a* abundance was ∼15% lower at 24 h compared to 6 h across all groups. Abundance of *crfbp1* (Fig. 6B; p_treatment_<0.001, p_time_<0.0001, p_treatment*time_<0.001) was ∼4-and ∼2.5-fold lower following cortisol treatment compared to vehicle- or cortisol+RU486-treated groups at 24 h (p<0.001 and p=0.004, respectively), but not at 6 h (both p>0.38). Additionally, levels of *crfbp1* in the cortisol+RU486-treated group were also ∼40% lower than the vehicle-treated group at 24 h (p=0.007), but not at 6 h (p=0.92). Abundance of *crfa1* (Fig. 6B; p_treatment_=0.77, p_time_<0.001, p_treatment*time_=0.01) was ∼25% lower in cortisol-treated spleens compared to vehicle-treated spleens at 6 h (p=0.04)—but not compared to cortisol+RU486-treated spleens (p=0.32)—and while levels at 24 h were ∼30-40% lower than at 6 h in vehicle-(p<0.001) and cortisol+RU486-treated groups (p=0.02), no change was detected in the cortisol treated group (p=0.69). Levels of *uts1a* (Fig. 6B; p_treatment_<0.001, p_time_<0.001, p_treatment*time_<0.001) were ∼30% and ∼130% higher in cortisol-treated spleens compared to vehicle-treated spleens at 6 h (p=0.03) and 24 h (p<0.001), respectively. However, while levels of *uts1a* in cortisol+RU486-treated spleens were not different from vehicle- or cortisol-treated spleens at 6 h (p=0.21 and p=0.64, respectively), cortisol+RU486-treated spleens had *uts1a* levels that were statistically intermediate to those of vehicle- (∼55% more) and cortisol-treated (∼30% less) spleens at 24 h (p<0.001 for both). Levels of *ucn3* (Fig. 6B; p_treatment_=0.30, p_time_<0.001, p_treatment*time_=0.31) were ∼50% lower at 24 h compared to at 6 h, but no treatment effects were detected.

## 4. Discussion

Interactions between stress-related hormones and the immune system have been well characterized in mammals, but these relationships remain unclear in non-mammalian vertebrates. We found that most components of the CRF system—which is an important regulator of immune responses in mammals (Dermitzaki et al. 2018; Zhu et al. 2011)—were expressed in both the spleen and head kidney of rainbow trout. Furthermore, we found that either *in vivo* or *in vitro* activation of splenic inflammatory responses caused transcriptional changes consistent with reduced CRFR1 activity and increased CRFR2 activity. Additional *in vitro* experiments suggest that immune-related changes in splenic CRFR1 and CRFR2 activity may be regulated by cortisol and NF-κB, respectively. Together, these data shed valuable light on interactions between immune responses and the CRF system in teleost fishes.

Consistent with the activation of an inflammatory response following vaccination, splenic transcript levels of proinflammatory cytokines (*il1b*, *il6*, *tnfa*, and *ifny*)—as well as protein levels of IL-1β and IFNγ—displayed marked increases (up to 3700-fold) in abundance within 6 h of vaccination, with peak induction of anti-inflammatory pathways occurring shortly thereafter (e.g., *il10* levels peaked at 24 h). We also observed changes in blood respiratory burst and neutrophil concentrations supporting a systemic immune response (Culbert et al., 2026 preprint). In parallel with these changes, we observed transcriptional evidence suggesting activation of CRFR2 in the period immediately following vaccination (i.e., elevated levels of *crfr2a* and *crfr2b* at 24 h post-vaccination). Additionally, levels of the CRFR2-specific ligand *ucn3* (Hsu and Hsueh, 2001; Manuel et al., 2014) were elevated at 6 h post-vaccination. In contrast, transcriptional changes suggest a suppression of CRFR1 activity since levels of *crfr1b* (with a similar, but non-significant reduction in *crfr1a* abundance), as well as *crfa1* and *uts1a* [ligands with high affinity for CRFR1 (Hosono et al., 2015; Manuel et al., 2014; Pohl et al., 2001)], were reduced following vaccination. Increased abundance of *crfbp1*—which can prevent UTS1 (and probably CRFa) from binding to CRFRs (Manuel et al., 2014)—at 6 and 24 h post-vaccination provides additional support for reductions in CRFR1 activation. Indeed, similar transcriptional changes supporting suppression of CRFR1 activity following vaccination—higher *crfbp* combined with lower *crfr1* and *crfa*—were also observed in the intestine of these same fish (Culbert et al., 2026 preprint). These contrasting patterns between CRFR subtypes suggest that they serve different functions and/or are located on different cell types within the spleen; both of which are supported by mammalian studies. For instance, immune-related increases in CRFR1 expression in splenic neutrophils contribute to IL-1β production in mice (Radulovic et al., 1999), whereas immune-related increases in the abundance of CRFR2 in murine splenic B cells (which do not express CRFR1) stimulate B cell apoptosis (Harlé et al., 2018). Additional support for divergent immune-related functions of the two CRFR subtypes in mammals also comes from murine mesenteric lymph node dendritic cells, where activation of CRFR1 (stimulatory) versus CRFR2 (inhibitory) had opposing effects on T cell proliferation and major histocompatibility complex 2 expression (Meng et al., 2015). Far fewer investigations of interactions between immune and CRF systems have been conducted in teleosts, and the only previous study to evaluate immune-related roles of the splenic CRF system in teleosts reported that reductions in phagocytosis following UTS1 treatment of phagocytes were mediated by CRFR1 in snakehead (Singh and Rai, 2011). In contrast, UTS1 increased rates of phagocytosis and suppressed production of pro-inflammatory cytokines in cultured head kidney monocytes/macrophages via an unknown CRFR subtype in ayu (Jiang et al., 2021). Additionally, CRF enhanced macrophage migration towards a wound site through CRFR1 and promoted a pro-inflammatory phenotype in larval zebrafish (van Heijningen et al. 2025). Overall, the immunoregulatory functions(s) mediated by the teleost CRF system likely vary between species and/or across tissues, highlighting the need for additional work to determine possible explanations for the CRFR-dependent responses to vaccination observed in the current study.

To further evaluate relationships between the CRF and immune systems—and elucidate potential mechanisms responsible for these relationships—we used *in vitro* spleen explant cultures. Based on transcriptional changes in all measured cytokines, spleen explants quickly (within 6 h) mounted a large inflammatory response that persisted for at least 24 h. Consequently, these preparations provided an excellent means of determining whether vaccine-related transcriptional changes in CRF system components were directly mediated by endogenous inflammatory responses occurring within the spleen (as opposed to being regulated by systemic factors originating from other tissues). Indeed, changes in transcript levels of *crfr2b*, *crfr1b*, *crfbp1*, *crfa1*, and *ucn3* in cultured spleen explants mirrored responses observed following vaccination (Table 1). Consequently, local inflammatory responses appear to be the primary immune-related regulator of most changes in the splenic CRF system. Curiously, two components (*crfr2a* and *uts1a*) showed opposing *in vivo* and *in vitro* responses, which may reflect the lack of regulatory contributions by endocrine factors during tissue culture (Castillo et al., 2009; Castro et al., 2011). Regardless, since transcriptional responses of most CRF system components in our *in vitro* preparations were comparable to vaccine-related responses, these *in vitro* preparations allowed us to evaluate which signalling pathways contribute to immune-related changes in the splenic CRF system.

Many immune-related responses—such as increased transcription of proinflammatory cytokines, including *il1b*, *il6*, *ifny*, and *tnfa* (Fenimore and Young, 2016; Liu et al., 2017)—are directly mediated by the activation of signalling pathways downstream of the transcription factor NF-κB. Indeed, we found that treatment of spleen explants with PDTC (an inhibitor of NF-κB) attenuated the increase in *il1b*, *il6*, and *ifny* transcript abundance that was observed in vehicle-treated explants. Levels of *il10* were also reduced, although, these effects were likely indirect since NF-κB does not directly regulate IL-10 (Bondeson et al., 1999). However, we did not detect any effect of PDTC treatment on *tnfa* transcription. The lack of effect in the current study could reflect the presence of additional TNFa paralogs in teleost fishes (Hong et al., 2013), which may have undergone sub- and/or neo-functionalization. Additionally, the upregulation of *tnfa* during culture (∼30-fold increase at 24h) was lower than that of either *il1b* (∼850-fold increase at 24h) or *il6* (∼650-fold increase at 24h), leaving comparatively less scope to detect PDTC-related effects. In addition to effects on cytokines, we found that NF-κB inhibition altered transcriptional responses of CRF system components. Specifically, we found that PDTC treatment decreased levels of *crfr2b* and *ucn3*, indicating reduced CRFR2 activity. Therefore, the increase in CRFR2 activity following vaccination (i.e., greater transcript abundance of *crfr2b* and *ucn3*) may reflect the presence of NF-κB responsive elements in the promoter region of these genes, as previously reported in the human CRF and CRFBP gene promoters (Behan et al. 1993; Want et al. 2012). Alternatively, their transcription may be regulated by another product that is controlled by NF-κB. However, since PDTC and cortisol treatment had similar effects on most cytokines measured in the current study but exerted different effects on the CRF system (Table 1), it is unlikely that changes in CRFR2 activity were mediated by any of the targeted cytokines. Clearly, future studies are needed to determine the mechanism(s) behind NF-κB’s effects on CRFR2 activity.

Given the important immunoregulatory functions of corticosteroids (Cain and Cidlowski, 2017; Turnbull and Rivier, 1999), as well as the established relationship between corticosteroids and the CRF system via the HPA/I axis (Aguilera, 1998; Best et al., 2024), we hypothesized that cortisol would be a major regulator of splenic CRF system activity. Indeed, in conjunction with the anti-inflammatory effects observed following cortisol treatment (e.g., reduced abundance of *il1b*, *il6*, *tnfa*, and *ifny* combined with increased abundance of *il10*), we also found that cortisol treatment increased the abundance of some CRF system components (e.g., *crfr2b*, *crfr1b*, and *uts1a*) while reducing levels of others (e.g., *crfr1a*, *crfbp1*, and *crfa1*). Most of these effects were deemed to be mediated (at least partially) by glucocorticoid receptor activation since co-treatment with a glucocorticoid receptor antagonist either suppressed or completely abolished effects of cortisol. However, while a few of these cortisol-mediated effects (e.g., increased *crfr2b* and reduced *crfa1*) matched what was observed following vaccination, many components showed opposing responses (e.g., increased *crfbp1* and reduced *crfr1b* and *uts1a* following vaccination). Since cortisol levels were only mildly elevated in vaccinated fish compared to saline-injected fish, and levels in both groups were consistent with cortisol concentrations previously reported for unstressed salmonids [e.g., ≤ 10 ng mL^-1^ (Culbert and Gilmour, 2016; Culbert et al., 2025b; Gamperl et al., 1994)], it is unlikely that the effects observed following vaccination were driven by changes in cortisol signalling. Additionally, while cortisol can clearly contribute to the regulation of splenic CRF system activity, these effects do not appear to be mediated by reductions in NF-κB signalling—despite this being one of the major mechanisms responsible for anti-inflammatory effects of corticosteroids in mammals (Ray and Prefontaine, 1994). Unlike the shared transcriptional responses of most cytokines following cortisol or PDTC treatment (suggesting that these cortisol-mediated effects may involve inhibition of NF-κB signalling), only a single component of the CRF system was similarly regulated by these two pathways (Table 1; increased *crfr2b* abundance). Furthermore, while inhibition of NF-κB signalling affected CRF system components associated with CRFR2 activity (i.e., *crfr2b* and *ucn3*), cortisol treatment primarily affected components associated with CRFR1 activity (i.e., *crfa1*, *uts1a*, *crfr1a*, *crfr1b*, and *crfbp1*; but also *crfr2b*). Therefore, these results likely reflect direct effects of corticosteroid receptor activation on CRF system components [i.e., glucocorticoid response elements are present in their promoter; (Huising et al., 2011; Malkoski et al., 1997)] and/or are related to changes in the activity of other corticosteroid-influenced signalling pathways, such as activator protein 1 or cAMP response element binding proteins (Cain and Cidlowski, 2017; McClennen and Seasholtz, 1999).

Interestingly, while many of the cortisol-mediated effects on the spleen CRF system observed in the current study are consistent with cortisol-mediated effects in Atlantic salmon (*Salmo salar*) gill filaments [e.g., upregulation of *crfr2b*, and downregulation of *crfr1a*, *crfbp2*, and *crfa1*; (Culbert et al., 2025a)], we observed contrasting responses in levels of *crfr1b* (i.e., downregulated in gill filaments and upregulated in the spleen). It is possible that this difference simply reflects species-specific transcriptional regulation of *crfr1b*, but it is also possible that the relationship between *crfr1b* and cortisol varies across salmonid tissues. Indeed, corticosteroids have contrasting transcriptional effects on CRF system components in peripheral tissues (Huising et al., 2011) and between different areas of the brain in mammals (Makino et al., 1994; Makino et al., 1995; Robinson et al., 1988), and a similar regulatory relationship is also slowly emerging in the brain of teleost fishes (Best et al., 2024). Consequently, future studies are needed to better understand the complex interactions between cortisol and different components of the CRF system in teleosts.

While CRFb is the primary CRF ligand within the teleost central nervous system (Bernier et al., 2009; Best et al., 2024; Maugars et al., 2022), our results show that neither CRFb paralog (CRFb1 or b2) is abundant in the spleen or head kidney. These data are consistent with our previous work reporting low abundance of CRFb outside of the central nervous system in salmonids (Culbert et al., 2022; Culbert et al., 2025a; Culbert et al., 2025b; Culbert et al., 2025c), indicating that auto- and paracrine contributions of CRFb within peripheral tissues (including the head kidney and spleen) are likely minimal. In general, CRF ligands were not abundant in the spleen (or head kidney), and while it is likely that the abundance of CRF system components (including ligands) is enriched in certain cell types versus others [e.g., B cells, macrophages, and/or neutrophils (Harlé et al., 2018; Radulovic et al., 1999; van Heijningen et al. 2025; Webster et al., 1990)], this suggests that circulating CRF peptides may have important regulatory roles in immune tissues. The primary source of circulating CRF peptides in teleosts is the caudal neurosecretory system [CNSS; (Bern et al., 1985; Rousseau et al., 2025; Winter et al., 2000)], which synthesizes large amounts of CRFb and UTS1 and releases them into circulation (Craig et al., 2005; Culbert et al., 2025b; Lu et al., 2004; Rousseau et al., 2025). We are not aware of any study which has evaluated how circulating CRF peptide abundance and/or CRF peptide production in the CNSS change following an immune challenge; however, future studies evaluating immune-related responses in the CNSS will be important for understanding the regulatory contributions of the CRF system during immune challenges.

Collectively, our findings indicate that the splenic CRF system in fish contributes to the regulation of inflammatory responses. Further research is necessary to identify the specific target(s) and function(s) of the splenic CRF system, as well as elucidate specific functions for CRFR1 and CRFR2. Determination of the immune functions served by the splenic CRF system in fish will provide novel insights into the spleen’s role in maintaining immunity and how dysfunction of these processes contributes to stress-related diseases.

## CRediT Authorship Contribution Statement

**Brett M. Culbert**: Conceptualization, Data curation, Formal analysis, Investigation, Methodology, Supervision, Validation, Visualization, Writing – original draft, Writing – review & editing. **Leah Grosman**: Investigation, Writing – review & editing. **Tania Rodriguez-Ramos**: Conceptualization, Investigation, Methodology, Validation, Writing – review & editing. **Brian Dixon**: Conceptualization, Funding acquisition, Methodology, Resources, Writing – review & editing. **Nicholas J. Bernier**: Conceptualization, Funding acquisition, Investigation, Methodology, Resources, Supervision, Writing – review & editing.

## Data Availability Statement

Data are deposited on Mendeley Data (doi: 10.17632/rvxspvpyzr.1).

## Acknowledgements

We would like to thank Marcia Chiasson and the staff at the Ontario Aquaculture Research Centre for providing us with the rainbow trout, as well as the fish husbandry assistance that was provided by Matt Cornish and Mike Davies at the Hagen Aqualab. Finally, we thank Carol Best for her assistance during the vaccine time course experiment.

