## Supplemental Table 1 for "Transcriptional regulation of the rainbow trout spleen corticotropin-releasing factor system in response to inflammatory challenges: roles of NF-κB and cortisol"

& Nicholas J. Bernier^1^

*^1^ Department of Integrative Biology, University of Guelph, Guelph, Ontario, Canada*

*^2^ Department of Biology, University of Waterloo, Waterloo, Ontario, Canada*

^†^Current Address: *Department of Biological Sciences, University of Cincinnati, Cincinnati, Ohio, USA*

**Supplemental Table 1.** Gene-specific primers used for real-time qPCR.

|  | Primer Sequence  (5’ to 3’) | Amplicon  Size (bp) | Efficiency  (%) | Accession  Number |
| --- | --- | --- | --- | --- |
| *crfa1* | F: CAAAATTCGCTCCAGTCCTC | 95 | 100 | XM_021613200 |
|  | R: TGCGTGAGCTGAAGTTGTAA |  |  |  |
| *crfa2* | F: ATTCGCCCCAATCTTCATT | 79 | 102 | XM_021589550 |
|  | R: TGAAGTAAAGCCCTGTTGACC |  |  |  |
| *crfb1* | F: CTGAAGGCATGTACCCAGAG | 110 | 102 | NM_001124286 |
|  | R: GACGAACCGGCTGATGTT |  |  |  |
| *crfb2* | F: GAAAGTCATGCACCCCAAG | 117 | 102 | NM_001124627 |
|  | R: AAGCCGCTGATGAACCTATT |  |  |  |
| *crfbp1* | F: CCAACACGTTGTCTATCTAC | 107 | 100 | NM_001124631 |
|  | R: TTTTATGGCAACCTTCAGGCT |  |  |  |
| *crfbp2* | F: CTCCCTATCTATCTCTCCAC | 122 | 100 | XM_036977080 |
|  | R: GTTATGGCGATCCTCAGGAT |  |  |  |
| *crfr1a* | F: TCCCTCGGAGACAGGAGTGT | 121 | 102 | XM_036940770 |
|  | R: CTTTTACGATGTGCGACTGGT |  |  |  |
| *crfr1b* | F: CGGCAATTTATAGGCTGTGTT | 114 | 99 | XM_021624572 |
|  | R: CCCACATGAACAACCAACAG |  |  |  |
| *crfr2a* | F: CACTGTACTGTACTAGTGGT | 83 | 100 | XM_021613636 |
|  | R: TCCTGCTTAAATGATCTCTGC |  |  |  |
| *crfr2b* | F: TAAATTCAGAACAGATGGGC | 82 | 100 | XM_021589160 |
|  | R: CATTGCCAGTAAGACTCTGAC |  |  |  |
| *ef1α* | F: CCATTGACATTTCTCTGTGGAAGT | 106 | 96 | NM_001124339 |
|  | R: GAGGTACCAGTGATCATGTTCTTGA |  |  |  |
| *ifny* | F: ACCCTGTTTTCCCCAAGGAC | 61 | 92 | NM_001124620 |
|  | R: ACCACACTCATCAACACCCTC |  |  |  |
| *il1b* | F: TGAGAACAAGTGCTGGGTCC | 148 | 107 | NM_001124347 |
|  | R: GGCTACAGGTCTGGCTTCAG |  |  |  |
| *il6* | F: AGGAGTTTCAGAAGCCCGTG | 51 | 104 | NM_001124657 |
|  | R: CCTGGTGCTGTGAGAACGAT |  |  |  |
| *il10* | F: CCGCCATGAACAACAGAACA | 105 | 93 | NM_001245099 |
|  | R: TCCTGCATTGGACGATCTCT |  |  |  |
| *rpl13a* | F: GGACAAGCTGCACTGGAGAG | 113 | 100 | XM_021574753 |
|  | R: GTGGGCTTCAGACGGACAAT |  |  |  |
| *tnfa* | F: CACACTGGGCTCTTCTTCGT | 155 | 98 | NM_001124374 |
|  | R: CAAACTGACCTTACCCCGCT |  |  | NM_001124357 |
| *ucn2a* | F: CATGCGTATGTGTCTGAGCC | 61 | 101 | CDQ61544 |
|  | R: TCTTCTCCCATCCTTTGGCAC |  |  |  |
| *ucn2b* | F: ACTCCTAAGTTGTGACATTGGA | 68 | 99 | XM_021570508 |
|  | R: CCAGACACAGGCTTAGATG |  |  |  |
| *ucn3* | F: GCACGGAACTTGGACATCAT | 91 | 100 | XM_021596023 |
|  | R: GTCTGAGACAGAGGCTGGAG |  |  | XM_036945073 |
| *uts1a* | F: GTTGCTGAAGGCTGGTGATA | 82 | 100 | NM_001124343 |
|  | R: CAACCGGGGATTTCTTTG |  |  |  |
| *uts1b* | F: GGCGACGCAGTTTCCTAC | 60 | 99 | XM_021612039 |
|  | R: GGGATTTCTCTGCAAATAACG |  |  |  |

*crf*, corticotropin-releasing factor; *crfbp*, corticotropin-releasing factor binding protein; *crfr*, corticotropin-releasing factor receptor; *ef1α*, elongation factor 1α; *ifny* interferon γ; *il*, interleukin; *rpl13a*, ribosomal protein L13a; *tnfa*, tumor necrotic factor α; *ucn*, urocortin; *uts*, urotensin.
